# Relating sparse/predictive coding to divisive normalization

**DOI:** 10.1101/2023.06.08.544285

**Authors:** Yanbo Lian, Anthony N. Burkitt

## Abstract

Sparse coding, predictive coding and divisive normalization have each been found to be principles that underlie the function of neural circuits in many parts of the brain, supported by substantial experimental evidence. However, the connections between these related principles are still poorly understood. Sparse coding and predictive coding can be reconciled into a learning framework with predictive structure and sparse responses, termed as sparse/predictive coding. However, how sparse/predictive coding (a learning model) is connected with divisive normalization (not a learning model) is still not well investigated. In this paper, we show how sparse coding, predictive coding, and divisive normalization can be described within a unified framework, and illustrate this explicitly within the context of a two-layer neural learning model of sparse/predictive coding. This two-layer model is constructed in a way that implements sparse coding with a network structure that is constructed by implementing predictive coding. We demonstrate how a homeostatic function that regulates neural responses in the model can shape the nonlinearity of neural responses in a way that replicates different forms of divisive normalization. Simulations show that the model can learn simple cells in the primary visual cortex with the property of contrast saturation, which has previously been explained by divisive normalization. In summary, the study demonstrates that the three principles of sparse coding, predictive coding, and divisive normalization can be connected to provide a learning framework based on biophysical properties, such as Hebbian learning and homeostasis, and this framework incorporates both learning and more diverse response nonlinearities observed experimentally. This framework has the potential to also be used to explain how the brain learns to integrate input from different sensory modalities.

**Author Summary:** Computational principles are often proposed to reveal the neural computations underlying brain functions. In the past three decades, sparse coding, predictive coding and divisive normalization have been three influential computational principles that have much success in different areas of neuroscience. Sparse coding offers insights into how the brain learns meaningful associations based on the hypothesis of brain being very efficient. With an emphasis on prediction, predictive coding provides an appealing hierarchical framework of only sending prediction errors to higher layers. Divisive normalization is a mathematical equation designed to account for the extensive nonlinearities in the brain. All these three computational principles along their variants have greatly improved our understanding of the underlying mechanism of the brain. Though connection between sparse and predictive coding has been studied previously, how sparse/predictive coding is connected to a seemingly different principle, divisive normalization, to provide a unified understanding of the brain is still unclear. In this paper, we show that sparse coding, predictive coding and divisive normalization can be connected from first principles. We propose a learning framework that is based on the hypothesis of efficiency, implemented with a predictive structure and displays response nonlinearities of divisive normalization. This framework can be potentially examined and used in a broader context such as multi-sensory integration.

## 1 Introduction

The structure of the brain’s networks enables it to perform complex daily tasks. Numerous experimental studies of the visual system, as well as other areas of the brain, have been conducted to uncover the underlying network structures, such as those responsible for simple cells and complex cells in cat’s primary visual cortex (V1) (Hubel and Wiesel, 1959, 1962, 1968). Different models have been proposed to account for various features of the experimental data, and this has resulted in a number of proposed neuronal models capable of describing some of the underlying mechanisms of the brain’s networks. Three such important models of network function are: (i) divisive normalization (Heeger, 1992), (ii) sparse coding (Olshausen and Field, 1996, 1997), and (iii) predictive coding (Rao and Ballard, 1999), each of which has had a significant impact due to their success in accounting for experimental data and providing specific mechanisms for the way in which the neural circuits function.

Divisive normalization, originally proposed by Heeger (1992), is a mathematical model that normalizes the response of individual model units (representing neurons in the real neural circuits) by the division of population response of the whole neural network; i.e., that the response of each neuron not only depends on its own activity but also on the responses of other neurons in the network. Divisive normalization was introduced to explain how responses of cells observed in cat V1 can display significant nonlinearities that linear models failed to account for. Heeger (1992) showed that a simple mathematical model of divisive normalization can explain many nonlinear response properties of V1 cells, such as contrast saturation and contrast invariance. Due to the wide diversity of nonlinear structure introduced through divisive normalization with different model parameters, divisive normalization has been used to explain a wide range of physiological data, from the visual system (Heeger, 1992) to the olfactory system (Olsen et al., 2010) and also multisensory integration (Ohshiro et al., 2011, 2017). Furthermore, divisive normalization has recently been integrated into deep neural networks, which improves the performance of image recognition (Miller et al., 2022) and image segmentation (Hernández-Cámara et al., 2023). For a review of divisive normalization see Carandini and Heeger (2012).

Sparse coding, originally proposed by Olshausen and Field (1996, 1997), finds an efficient representation of the sensory input using a linear combination of a set of basis vectors. Sparse coding is based upon two important assumptions, namely that the model generates a linear reconstruction of the input and this linear reconstruction is efficient. The notion of efficiency adopted in sparse coding is based on the principle of redundancy reduction proposed by Barlow (1961, 1989). Olshausen and Field (1996, 1997) applied sparse coding to natural images and showed that this model can learn Gabor-like features after learning, which resemble the observed receptive fields (RFs) of simple cells in V1. An appealing feature of sparse coding is its learning ability, namely that the model describes how the connection weights can be learnt based on the statistics of the input visual data, in a way that is similar to synaptic plasticity. Apart from its success in learning RFs of cells in the visual cortex, sparse coding has also been successfully applied to explain experimental results in other brain areas, such as auditory cortex (Smith and Lewicki, 2006) and hippocampal formation (Lian and Burkitt, 2021, 2022; Lian et al., 2023). For a review see Beyeler et al. (2019).

Predictive coding, originally proposed by Rao and Ballard (1999), provides a hierarchical predictive framework that is inspired by the observation of the extensive feedback in the cortex. Predictive coding assumes a hierarchical network structure, where units in the lower layers of the hierarchy project to the successive higher layers via feedforward connections, while units in the higher layers project back to the lower layers via feedback connections between the adjacent layers. The residual error, defined as the difference between the feedforward input to a layer and the feedback from its adjacent higher layer, is computed and generates the response of the units in the lower layer that project to the upper layer. The feedback from higher layers of the hierarchy represents the prediction at that layer of the input, and thus the residual error is often referred to as *prediction error* in predictive coding. Note that this framework is predictive in a hierarchical sense rather than a temporal sense. The essence of predictive coding is that the higher layers in the network hierarchy constantly make prediction of the current inputs, compute the mismatch between input and prediction, and only transmit that mismatch (prediction error) to the higher layer. The notion of only transmitting mismatch to the higher layer is similar to the principle of efficiency in sparse coding. Rao and Ballard (1999) showed that predictive coding, when used to model the visual pathways of the brain, can learn simple-cell-like RFs and display some non-classical RF properties such as end-stopping. A Bayesian framework was later constructed using predictive coding, which provides an appealing model for many different brain areas (Knill and Pouget, 2004; Friston, 2010; Bastos et al., 2012).

Built upon these three significant neural mechanisms - sparse coding, predictive coding and divisive normalization - many computational and experimental studies have successfully been able to expand our understanding of particular brain functions. The connection between sparse coding and predictive coding has been studied previously (Boutin et al., 2020), but how they are related to divisive normalization towards a unified framework is still lacking. Though the direct connection between these three models may not be immediately apparent, we propose that these three models are closely related through the underlying network structures by which they are implemented. In this paper, we provide a framework that connects sparse coding, predictive coding and divisive normalization from first principles.

## 2 Materials and Methods

### 2.1 Sparse coding

Sparse coding finds an efficient representation of the sensory input using a linear combination of a set of basis vectors (Olshausen and Field, 1996, 1997). Denote

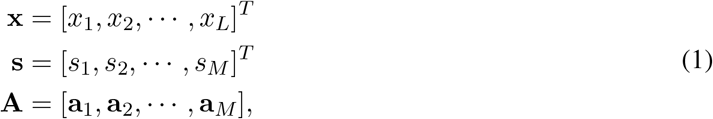

where **x** is a *L ×* 1 vector that represents the sensory input such as visual or auditory stimuli that has been pre-processed upstream, **s** is a *M ×* 1 vector that represents the response of *M* model units, and **A** is a *L × M* matrix that contains *M* basis vectors **a**_1_, **a**_2_, *···*, **a**_*M*_. Each basis vector, **a**_*i*_ (*i* = 1, 2, *···, M*) is a *L ×* 1 vector that represents one feature used to reconstruct the input such that the input, **x**, can be represented by a linear combination of basis vectors weighted by model responses:

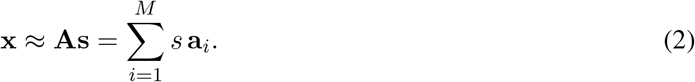

Sparse coding finds the linear representation (Eq. 2) using sparse responses, namely that only part of the model responses, *s*_*i*_ (*i* = 1, 2, *···, M*), will be active. This can be formulated as an optimization problem of minimizing the following cost function:

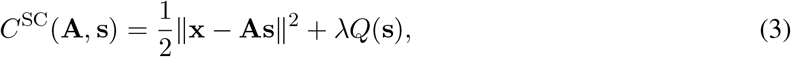

where 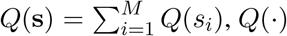 is a function that penalizes large responses, and *λ* is a constant parameter that scales the penalty function. In other words, the first term on the right-hand-side of Eq. 3 represents the reconstruction error and the second term determines the sparsity of model units. Therefore, by minimizing *C*^SC^(**A, s**), one finds the linear reconstruction of the input using sparse responses. However, there is always a trade-off between the reconstruction fidelity and the response sparsity. Note that Eq. 3 is the same as Eq. 2 in Olshausen and Field (1996) and Eq. 14 in Olshausen and Field (1997).

Assuming that *Q* (*·*) is differentiable and given **A**, the dynamics of **s** can be derived from taking the partial derivative of *C*^SC^(**A, s**):

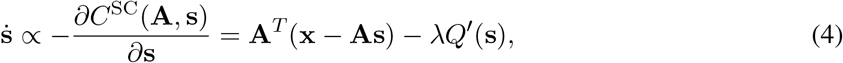

where *Q′* (*·*) represents the derivative of *Q*(*·*) and *Q*^*′*^ (**s**) = [*Q*^*′*^ (*s*_1_), *Q*^*′*^ (*s*_2_), *···, Q*^*′*^ (*s*_*M*_)]^*T*^. Eq. 4 describes how, given input **x**, an iterative procedure enables us to calculate the response of model units. The response **s** may or may not reach a stable equilibrium, depending on the properties of this dynamic system.

The learning rule for the connection weights **A** can also be derived by taking the partial derivative of *C*^SC^(**A, s**):

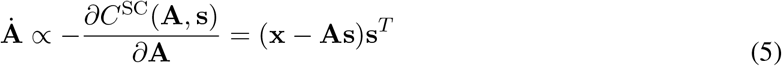

L2 norm of **A** is then adapted.

Generally, when simulating sparse coding models, the learning rule in Eq. 5 is applied once only after some iterations of computing the model response according to the model dynamics described by Eq. 4, with the assumptions that (i) the time scale of learning/plasticity is longer than that of model responses, and (ii) the model response iterates to some stable value. The learning rule in Eq. 5 may cause basis vectors in **A** to grow without bound, so L2 norm normalization of each vector, **a**_*i*_, is used in the original work of sparse coding (Olshausen and Field, 1997) and some subsequent studies (Zhu and Rozell, 2013; Lian et al., 2019).

### 2.2 Predictive coding

Predictive coding, originally proposed by Rao and Ballard (1999), is a hierarchical model in which prediction errors are propagated to the higher levels of the network hierarchy. Predictive coding can have multiple layers of the network, but in this study we only look at a two-layer predictive coding network model because it is the fundamental building block of multi-layer predictive coding and it can be explicitly shown how it is related to sparse coding and divisive normalization. Combining Eqs. 4 and 5 in Rao and Ballard (1999), the cost function of predictive coding can be written as

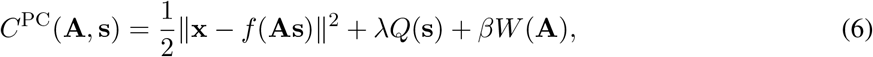

where *f* (*·*) can be interpreted as the neuronal activation function, *f* (**As**) is the prediction of the input, *Q*(**s**) is the prior constraint on the model responses, and *W* (**A**) puts some regularization on the basis vectors that are the columns of **A**.

Similar to the sparse coding case, the model dynamics and learning rule of predictive coding can be derived by taking the corresponding partial derivatives of *C*^PC^(**A, s**):

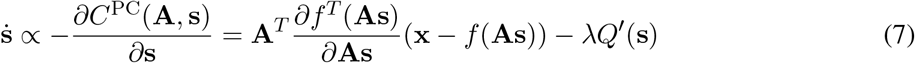

and

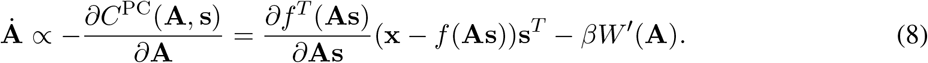

The learning rule of predictive coding is applied after some iterations of computing the model responses.

### 2.3 Divisive normalization

Divisive normalization was first proposed by Heeger (1992) to account for the nonlinear physiological responses of V1 cells. As reviewed by Carandini and Heeger (2012), one commonly used variant of divisive normalization is given by

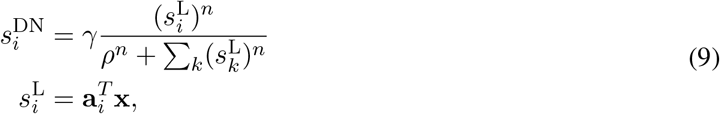

where **x** is the sensory input, **a**_*i*_ represents the synaptic weights connecting input with neuron *i* and is normally hand-wired, 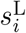 is the linear response, 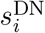 is the response of neuron *i* after divisive normalization, and *γ, ρ* and *n* are the parameters that determine the shape of the response curve of divisive normalization. Note that 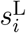 can also incorporate a rectifying function, but we use a linear function for the purpose of analytical analysis below. The model structure of divisive normalization is shown in Fig. 1.

**Figure 1:**
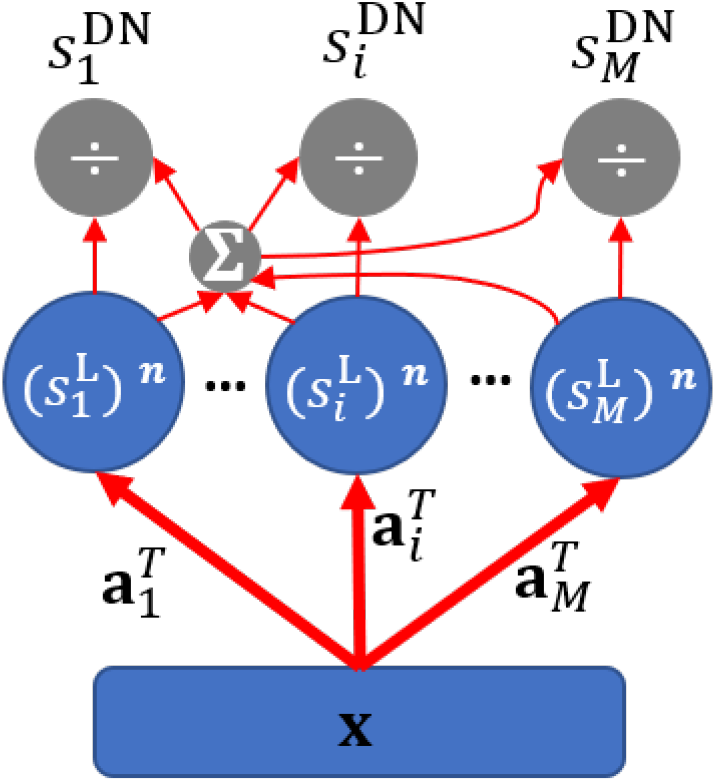
Model structure of divisive normalization (DN). The linear response is first generated; some form of nonlinearity (exponential function with exponent *n*) is then applied; next, the response of each unit is divided by the sum of responses of all the units in the network.

Divisive normalization introduces nonlinearity into the model through the calculation of the neural responses in Eq. 9, which also introduces competition between model units through the scaling of the response of each unit by the response of the whole population. Note that divisive normalization, similar to sparse coding and predictive coding, involves the definition of connection weights, but unlike sparse coding and predictive coding, the synaptic weights are not learnt through a learning process.

### 2.4 Applying the model on vision

In this paper, we propose a model based on sparse coding with the structure of predictive coding while incorporating the response nonlinearities of divisive normalization. We apply the proposed model on the context of visual processing in the brain. First, we train the model using natural images to learn V1 simple cells. Second, we investigate the response nonlinearity of learnt V1 simple cells by looking into their response vs. input relationship.

#### 2.4.1 Training the model with natural images

Similar to previous studies (Olshausen and Field, 1996, 1997), ten pre-processed natural images are used to train the model and image patches with size 16 *×* 16 sampled from these natural images are used as the input to the model. In each training epoch, a batch of 100 image patches are used to accelerate learning. Model dynamics and learning rule are implemented according to Eqs. 10 and 11 and model responses are kept non-negative.

#### 2.4.2 Investigating the response nonlinearity of learnt model

After training, we investigate response vs. input relationship of learnt model cells by presenting sinusoidal gratings with different levels of contrasts. First, we only include model cells to plot their response vs. input in response to optimal sinusoidal gratings if they meet two requirements: a) the receptive field of the model cell can be well fitted by a Gabor filter with fitting error smaller than 40%; b) the receptive field is located in the central region and well within the visual field. Second, we present optimal sinusoidal gratings at a contrast level with the preferred spatial frequency and preferred orientation determined by the fitted Gabor filters, and a range of different spatial phases (0 *−* 360°) to mimic moving sinusoidal gratings presented to animals when investigating response properties of V1 cells in experimental studies. Third, the mean value of the stable response of the model cell to the above sinusoidal gratings across all spatial phases is then computed as the model response to the optimal sinusoidal gratings at a specific contrast level. Fourth, we vary the contrast level of the optimal sinusoidal gratings and collect the corresponding model responses. Lastly, we plot the response vs. contrast curve to investigate the response nonlinearity of the learnt model cell.

The response vs. contrast curve for suboptimal sinusoidal gratings can be obtained in the same manner except the sinusoidal gratings have a suboptimal orientation (e.g., 30° away from the preferred orientation).

The model was simulated in MATLAB (R2024b, USA) and the code is available at https://github.com/yanbolian/Relating-Sparse-Predictive-Coding-to-Divisive-Normalization.

## 3 Results

### 3.1 Relationship between sparse coding and predictive coding

In this section, we first demonstrate the similarity between sparse coding and predictive coding. Though the connection between these two models has been mentioned in a previous study (Boutin et al., 2020), we explicitly connect these two models from first principles and refer them to as sparse/predictive coding, which will be further connected with divisive normalization in later sections.

Predictive coding is well known to be closely related to sparse coding, or efficient coding more broadly speaking, because only the prediction error is fed to higher levels of the network hierarchy. Furthermore, it can be shown from first principles that sparse coding and predictive coding are identical in a two-layer neural network under some parameter choices.

In the original cost function of predictive coding (Eq. 6), the term *βW* (**A**) applies some penalties on the connection weights **A** during the learning process, since its derivative appears in the learning rule described by Eq. 8. Rao and Ballard (1999) used the sum of squared response in the cost function that regulates the L2 norm of learned basis vectors (**A**), which is essentially similar to L2 normalization used in sparse coding models during the learning process. Therefore, *− βW* (**A**) in Eq. 6 can be removed and *βW* (**A**) in Eq. 8 can be approximated by a L2 normalization. Additionally, in the case where *f* (*·*) is the linear function, *f* (*y*) = *y*, then the cost function (Eq. 6), model dynamics (Eq. 7) and learning rule (Eq. 8) of predictive coding become identical to those of sparse coding (Eqs. 3, 4 and 5). Note that *f* (*y*) = *y* was also used in end-stopping simulations to illustrate predictive coding in the original study by Rao and Ballard (1999). In addition, the learning rules of both sparse coding (Eq. 5) and predictive coding (Eq. 8) have a Hebbian component, namely (**x** *−* **As**)**s**^*T*^ for sparse coding and (**x** *− f* (**As**))**s**^*T*^ for predictive coding. By interpreting (**x** *−* **As**) or (**x** *− f* (**As**)) as the response of error neurons that compute the difference between input and model reconstruction, these learning rules can be seen to obey the Hebbian learning principle.

In summary, when the weight regularization term is replaced by L2 normalization and the feedback activation function, *f* (*·*), is linear, sparse coding is identical to predictive coding. A unified framework with model diagram and equations is given below.

The exposition above shows how sparse coding and predictive coding can be described in the same framework, although their original formulation had different goals, namely sparse coding is about the sparsity in the network model while predictive coding is concerned with the goal of the resulting neural processing. Here a unified framework for sparse coding and predictive coding is illustrated by implementing the principle of sparse coding in a two-layer model with the structure of predictive coding. We refer to this model as sparse/predictive coding (SPC) model, similar to Boutin et al. (2020). The structure of the SPC model is shown in Fig. 2.

**Figure 2:**
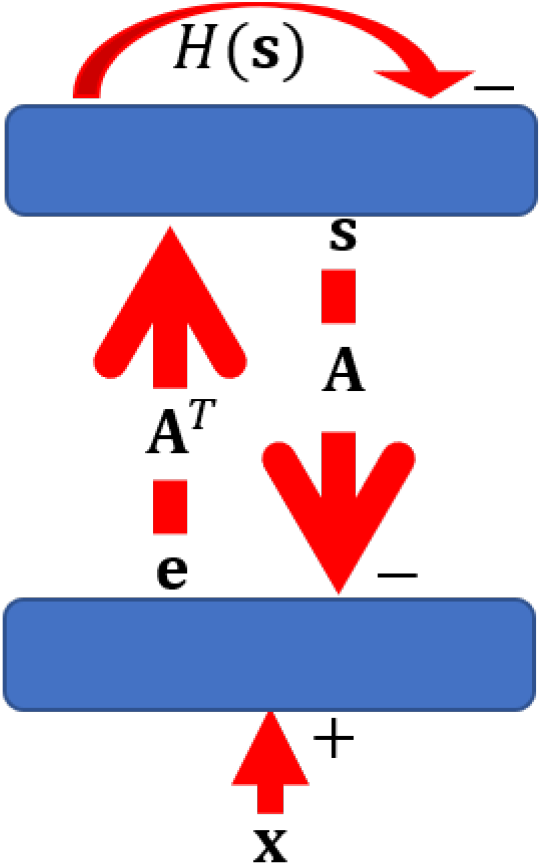
Model structure of sparse/predictive coding (SPC). The first layer computes the prediction error between the input and reconstruction of the model, and then sends error signals to the second layer. The second layer takes input from the first layer and incorporates the response-regulating mechanism, *H*(*·*). This two-layer network implements the dynamics described in Eq. 10.

The similarity between sparse coding (Eq. 4) and predictive coding (Eq. 7) enables the dynamics and learning rule of the SPC model to be described as a stage of *response iteration*:

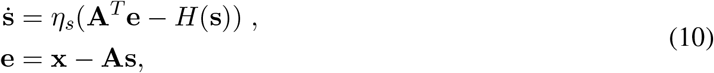

followed by the *learning step*:

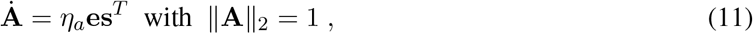

where **e** is the prediction error, *H*(**s**) = *λQ* (**s**) = [*H*(*s*_1_), *H*(*s*_1_), *···, H*(*s*_*M*_)]^*T*^ is the mechanism that regulates the responses of model units, *η*_*s*_ is the update rate when computing the responses, *η*_*a*_ is the learning rate of updating connection weights, and the magnitude of **A** is regulated by weight normalization (**A** _2_ = 1) at the end of each learning step, similar to previous studies (Olshausen and Field, 1997; Rozell et al., 2008; Zhu and Rozell, 2013; Lian et al., 2019).

The brain has self-regulating mechanisms called homeostatic plasticity that stabilizes the neural system via synaptic scaling and response adaptation (Turrigiano, 2011, 2017). Sparse coding and predictive coding have components that are similar to homeostasis. The weight normalization in sparse coding (Eq. 5) and weight regularization in predictive coding (Eq. 8) scale weights so that the learning process is stable, playing a role similar to homeostatic synaptic scaling. In the cost functions of sparse coding (Eq. 3) and predictive coding (Eq. 6), *Q*(*·*) is the sparsity term that promotes sparse responses of the model units and its derivative is denoted as *H*(*·*), as appeared in the SPC model (Eq. 10). Consequently, *H*(*·*) can be understood as a homeostasis function that self-regulates individual neuronal responses and should only depend on the homeostatic mechanism of individual neurons.

The model responses are computed in response iteration using Eq. 10 and then the learning rule of SPC is applied in the learning step using Eq. 11. Note that Eq. 11 is a local Hebbian learning rule in this model structure. *η*_*s*_ needs to be chosen carefully to ensure reasonable convergence of the model responses (see Fig. 4C for an example). *η*_*a*_ also needs to be much larger than *η*_*s*_ (so that the weight dynamics is on a much slower timescale than the response dynamics) and to be chosen such that **A** can learn meaningful structure at a reasonable speed.

When this SPC model is related to the neural circuits of the brain, neurons in the first layer compute the prediction error and feed the error signals to the second layer, where the responses are computed according to the dynamics in Eq. 10. **A**^*T*^ and **A** can be interpreted as the feedforward and feedback synaptic connections between two layers. This structure was used by Lian et al. (2019) to implement sparse coding of V1 simple cells with separate ON and OFF LGN channels and the learnt receptive fields, given by **A**^*T*^, agree well with experimental data and can account for many experimental phenomena observed in the LGN-V1 pathway.

An alternative way of implementing sparse coding in a feedforward network is through inhibitory lateral connections, which promote competition that eventually leads to sparse responses (Rozell et al., 2008; Zylberberg et al., 2011; Brito and Gerstner, 2016), and the role of the lateral connections is to decorrelate responses across the network, so that the model can learn different features. However, the mathematical equations underlying different implementations of sparse coding are essentially the same. Therefore, the framework of SPC proposed here can unify a range of different models related with sparse coding and predictive coding.

### 3.2 The homeostasis function in sparse/predictive coding determines response nonlinearity

The dynamics of SPC model described in Eq. 10 can be reformulated into one equation:

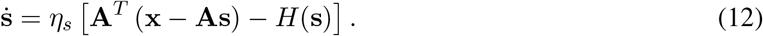

The equilibrium, **s**^EQ^, of this differential equation is reached when 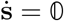, namely when

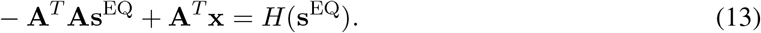

Solutions to Eq. 13 are the steady state response when the iterative process of solving Eq. 12 converges. If we define *Y* (**s**) as a linear function of **s** with form *Y* (**s**) := *−* **A**^*T*^ **As** + **A**^*T*^ **x**, where *−* **AA**^*T*^ is the slope and **A**^*T*^ **x** is the offset of *Y* (**s**), then the solutions **s**^EQ^ to Eq. 13 are just the intersects between *Y* (**s**) and *H*(**s**). Function *H*(*·*) is fixed for any given neuron because it depends on the self-regulating homeostatic mechanisms of that particular neuron. As *Y* (**s**) shifts, corresponding to its offset **A**^*T*^ **x**, the intersect between these two functions determines the analytical equilibrium of the activity, **s**^EQ^, given the current input **x**. A simplified one-dimensional illustration is shown in Fig. 3.

**Figure 3:**
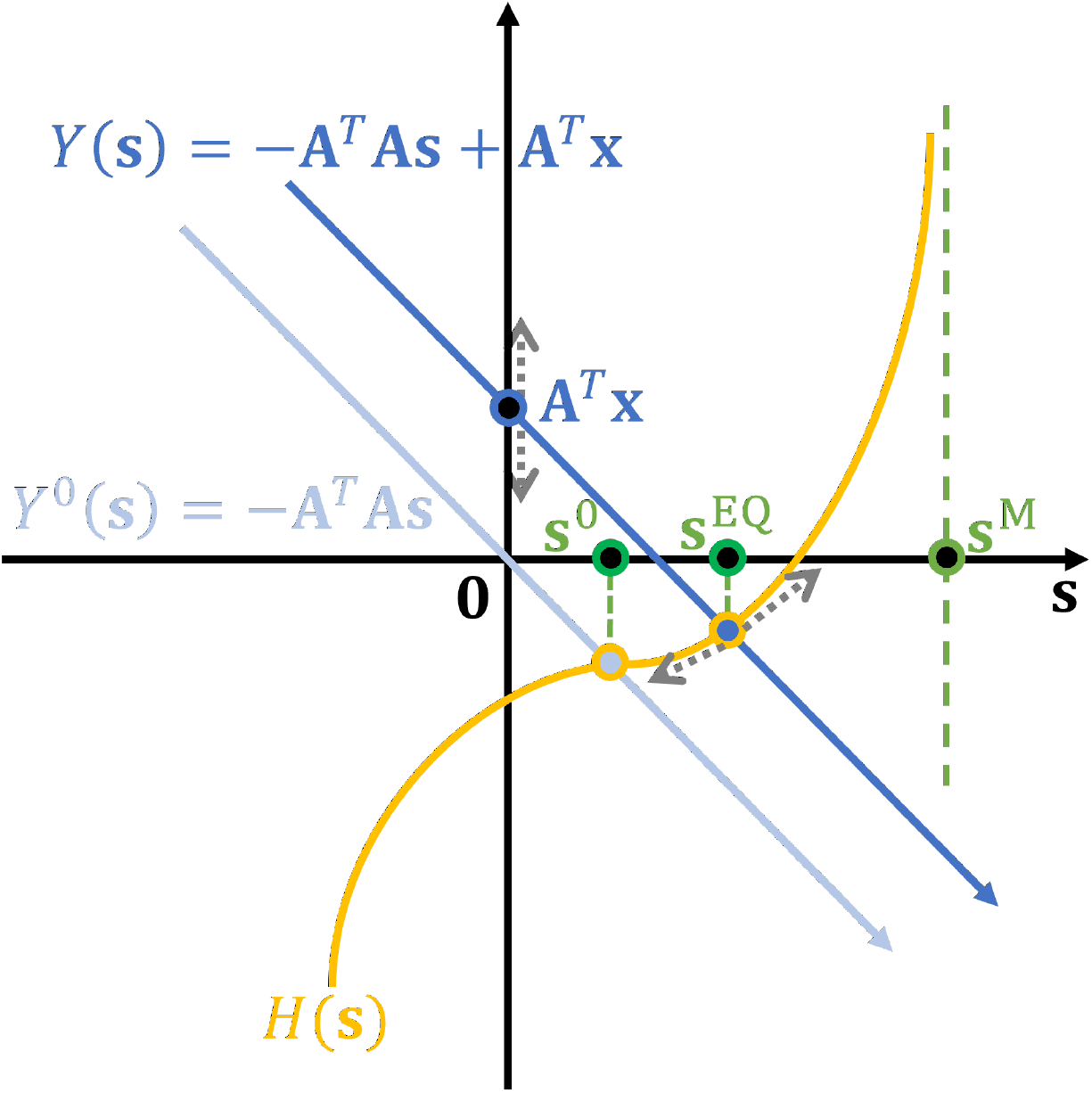
Equilibrium of sparse/predictive coding model explained in one-dimensional space. The intersect between *Y* (**s**) = *−* **A**^*T*^ **As** + **A**^*T*^ **x** and *H*(**s**) determines the model’s analytical equilibrium, **s**^EQ^. As input **x** changes, *Y* (**s**) shifts with the corresponding offset **A**^*T*^ **x** and the intersect moves along the curve of function *H*(**s**). When **x** = 0, the intersect is the model’s spontaneous response, **s**^0^. If *H*(**s**) has a vertical asymptote at **s** = **s**^M^, then **s**^M^ becomes the analytical upper bound upon response of the model.

**Figure 4:**
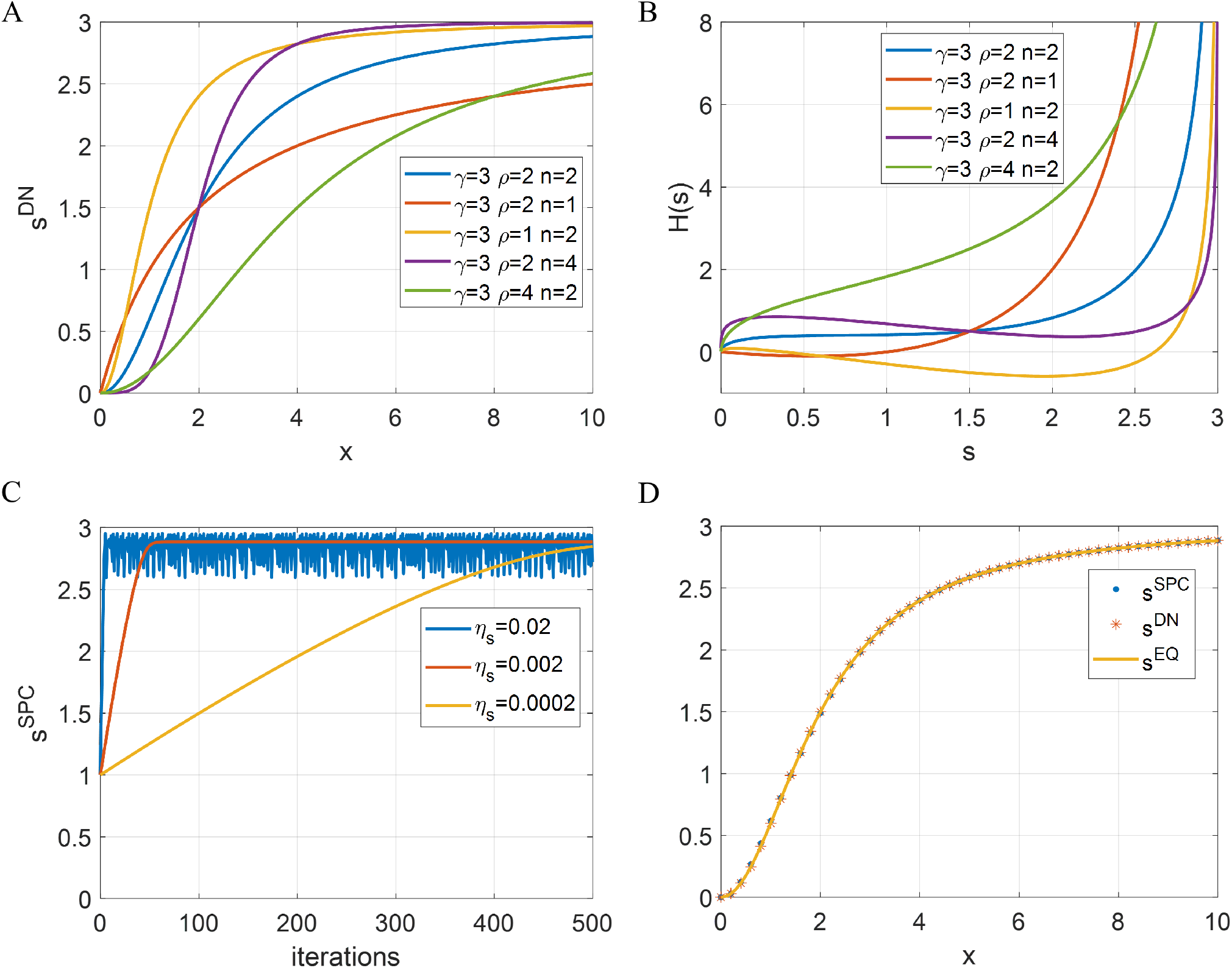
Divisive normalization (DN) can be identical to sparse/predictive coding (SPC) for single-neuron scenario. (A) DN described by Eq. 16 with different choices of parameter values. (B) Homeostasis function, *H*(*s*), of the equivalent SPC model, described by Eq.19. (C) Response of SPC vs. number of iterations for input, *x* = 10, with different values of *η*_*s*_ (inserted panel) when *γ* = 3, *ρ* = 2 and *n* = 2. (D) Responses of DN and SPC with *η*_*s*_ = 0.002 vs. different levels of input *x* when *γ* = 3, *ρ* = 2 and *n* = 2. The solid line is the analytical response vs. input curve that are solved numerically from *H*(*s*).

Consider now the case of the spontaneous activity of the neuron, namely the response when the input **x** is zero. When there is no external input to the network (i.e., **x** = 0), the spontaneous response of the SPC network is determined by the intersect between the curves *Y* (**s**) = *−* **A**^*T*^ **As** and *H*(**s**). Therefore, given **A**, the choice of *H*(*·*) determines the spontaneous response of the model. When input **x** changes linearly, *Y* (**s**) also changes linearly, so how the intersect **s**^EQ^ changes solely depends on the shape of *H*(**s**). In other words, *H*(*·*) determines the response nonlinearity of the model. For example, if *H*(**s**) is a linear function of **s**, model equilibrium is linearly dependant on the input **x**; if *H*(**s**) is highly nonlinear, there will be a great nonlinearity between model equilibrium and input; if *H*(**s**) approaches infinity with a vertical asymptote **s** = **s**^M^, then the model equilibrium will saturate at **s**^M^ although the input **x** keeps increasing. Given any homeostasis function *H*(*s*), one can obtain the analytical response vs. input curve by computing the intersection between *H*(*s*) and *Y* (*s*) while changing *s* linearly, which can be normally computed numerically. When there is no homeostasis namely that *H*(*s*) = 0, the x-axis intersect of function *Y* (**s**) will be the equilibrium of the model (see detailed analysis in the Appendix).

The Jacobian matrix of the model dynamic (Eq. 12) is also applied to analyze the stability of the system at the equilibrium, where

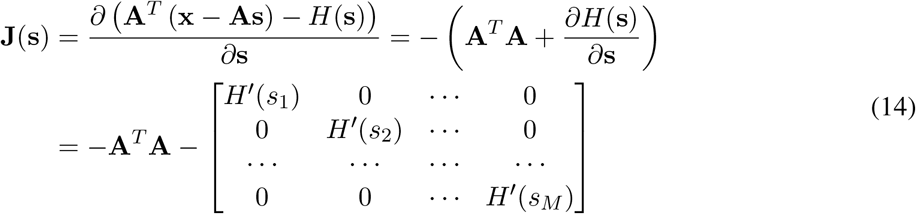

The eigenvalues of **J**(**s**) can be represented by

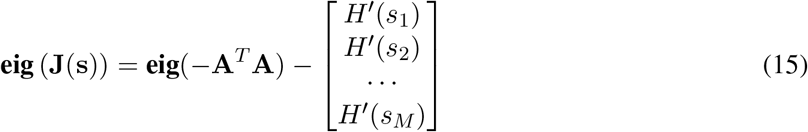

where **eig**(*·*) represents the vector of all eigenvalues of the matrix. Because *J* (**s**) = *−* **A**^*T*^ **A** is a negative definite matrix, the eigenvalues are all real and negative. If *H*(*·*) is monotonically increasing (i.e., *H*′ (*·*)*≥* 0), eigenvalues of **J**(**s**) will also be negative real values, indicating the stability of the equilibrium. When *H*(*·*) is not always monotonically increasing, the stability depends on the exact values of **eig** (**J**(**s**)).

### 3.3 Relationship between sparse/predictive coding and divisive normalization

Divisive normalization (DN) is fundamentally different from sparse/predictive Coding (SPC). Unlike SPC, DN is based upon a physiological model that was introduced to explain the nonlinear neural response properties in the brain, rather than a learning model with neural dynamics and synaptic plasticity. Specifically, for the model responses, SPC applies an iterative procedure described by Eq. 10, while the response of DN can be directly calculated by Eq. 9; for the learning rule, SPC incorporates learning and can describe how some brain properties can emerge via learning (Olshausen and Field, 1996, 1997; Zhu and Rozell, 2013; Lian et al., 2019), while DN has no learning rule and can only be used for explaining physiological neural response data. DN has a wide diversity of response nonlinearity with different parameters of the model, so it has been successful to account for the neural response nonlinearities found in experimental studies (Carandini and Heeger, 2012).

As demonstrated in the previous section (Section 3.2), the shape of the homeostasis function, *H*(*·*), determines the response nonlinearity of the SPC model, so SPC may potentially display the response nonlinearity of DN models, given a suitable choice of *H*(*·*).

#### 3.3.1 SPC and DN can be identical for a single-neuron model

Consider the case where there is only one neuron in the first layer and one neuron in the second layer for both the SPC and DN models, that the input is nonnegative, and the connection weight is 1. The DN model computes the response as

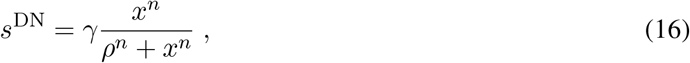

where *γ* determines the saturation value for the response, *ρ* determines the point where the response is *γ/*2, and *n* determines the curvature of the normalization curve. Note that the above equation has the form of Naka-Rushton function (Naka and Rushton, 1966) and has been used to account for response linearities in the retina and olfactory system (see Carandini and Heeger (2012) for a review). Some examples of DN described by Eq. 16 are shown in Fig. 4A.

For the SPC model, the equation that solves the analytical equilibrium of the output neuron (Eq. 13) becomes

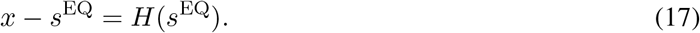

From Eq. 16 of DN, the solution for *x* is given by

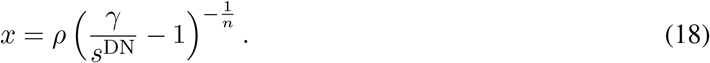

If DN and SPC are equivalent for this scenario (i.e., *s*^DN^ = *s*^EQ^), the homeostasis function in SPC can be derived from Eqs. 17 & 18:

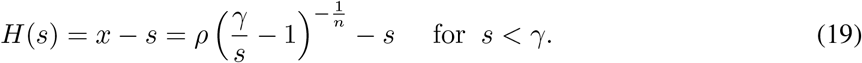

Some homeostasis functions of SPC described by Eq. 19 are shown in Fig. 4B. When *s* is small, *H*(*s*) has a small cost on responses, though it displays various shapes for different sets of parameters. As *s* approaches 3 (the value of *γ*), *H*(*s*) has a very high cost that prevents the response from exceeding the saturation. Therefore, for a single neuron scenario, SPC with a suitable choice of the homeostasis function can be equivalent to DN model that displays response nonlinearity and response saturation. Note that the homeostasis function (Eq. 19) is derived from Eq. 16 where *s*^DN^ is always smaller than the saturation value *γ*. Consequently, *H*(*s*) in Eq. 19 only applies for values *s < γ*.

The simplest scenario described above is implemented with *γ* = 3, *ρ* = 2 and *n* = 2. DN is implemented using Eq. 16. SPC is implemented by an iterative procedure described by Eq. 10 with *H*(*s*) given by Eq. 19 and 500 iterations of computing the response to any input. When the input is *x* = 10, the response of SPC model vs. number of iterations is plotted in Fig. 4C, which shows that the response of SPC gradually evolves to the equilibrium and its dynamics depends upon the value of *η*_*s*_. *s*^SP^ starts with a value of 1 because it is initialized to 1 prior to the iterative process. When computing the response of the SPC model, the value of *η*_*s*_ requires careful consideration. If *η*_*s*_ is too small, it takes more iterations to reach the equilibrium (e.g., yellow line in Fig. 4C). However, if *η*_*s*_ is too large, *s*^SP^ may become oscillatory (e.g., blue line in Fig. 4C). Additionally, if *s*^SP^ overshoots the saturation value, *γ*, during simulation of the response iteration (Eq. 10), this will cause the SPC model to break down because *H*(*s*), described by Eq. 19, is well-defined only for *s < γ*.

Fig. 4D shows that SPC and DN can generate equivalent results where the model response of SPC after convergence is identical to the model response of DN for all inputs *x ≥* 0. Furthermore, in this simplest scenario, the connection weight is 1, so *Y* (*s*) = *s* + *x* in Fig. 3 and the analytical response vs. input curve of the SPC model can be numerically solved by finding the intersect between *Y* (*s*) and *H*(*s*) while varying *x*. As shown in Fig. 4D, the analytical response curve closely matches the corresponding DN curve.

The analysis of this simple scenario illustrates that although SPC and DN are different in terms of their model dynamics and model structure, they can nevertheless generate equivalent network activity given a suitable choice of the homeostasis function in the SPC model. However, the homeostasis function in SPC enforces a rigid constraint upon the response, namely that the response has to be always smaller than *γ* during the iterative procedure of computing responses. When the network has more complexity, namely as the number of neurons increases, this assumption can be violated.

### 3.3 Relationship between SPC and DN in a multi-neuron network

If there are multiple neurons in the network, denote **x** = *α***x**_0_ where **x**_0_ is a normalized stimulus and *α* is a scalar that determines the strength of the input **x**. The divisive normalization definition, Eq. 9, then becomes

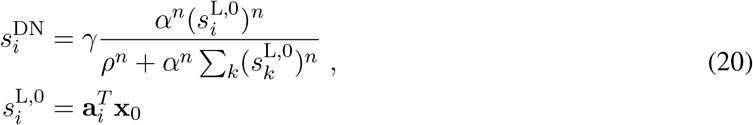

where 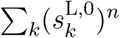 represents the population response to a normalized stimulus **x**_0_ with some nonlinearities determined by the exponent *n*. Denoting 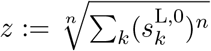, then Eq. 20 becomes

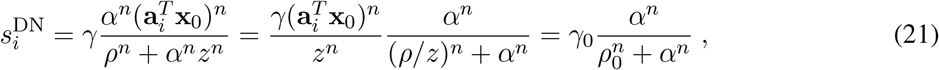

where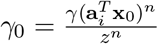. Comparison of Eq. 16 and Eq. 21 shows that the relationship between the model response of DN and the input magnitude *α* can be reduced to expression for the single-neuron scenario analyzed in previous section. Consequently, *H*(*·*) of SPC can be chosen to account for response nonlinearities described by DN. However, given its dependence on the weights **a**_*i*_ of other neurons, it is no longer straightforward to obtain a closed-form mathematical formula that provides the equivalent SPC model for any given DN model (as done for single-neuron case in Eq. 19).

### 3.4 Sparse/Predictive coding can learn Gabor-like simple cells with contrast saturation previously explained by divisive normalization

One notable difference between DN and SPC is the learning ability: DN is a model of only the physiological responses, and it can account for many nonlinear response properties, whereas the SPC model has a learning process; i.e., the SPC model, unlike the DN model, can learn important features of the input, given a suitable choice of the homeostasis function (Olshausen and Field, 1996, 1997; Rao and Ballard, 1999). As demonstrated by Zhu and Rozell (2013), sparse coding can be used to explain a number of nonlinear receptive field properties of V1 cells such as contrast invariance and end-stopping. However, contrast saturation of V1 cells, which is a classical nonlinear property (Ohzawa et al., 1982, 1985; Freeman et al., 2002), can not be captured in the sparse coding model of Zhu and Rozell (2013), whereas it can easily be captured by DN (Heeger, 1992). In a simple scenario in the section above, DN can be equivalent to SPC with suitable choice of the homeostasis function, *H*(*s*) (Eq. 19 and Fig. 4B), which suggests that SPC can account for saturation at least for a single-neuron scenario (Fig. 4D). Saturation in the SPC model is built into the homeostasis function, *H*(*s*) (Eq. 19), that has a very large penalty (infinity in this case) at the saturation point.

In this section, we show that SPC with a novel homeostasis function can learn Gabor-like simple cells as well as displaying the property of contrast saturation that is previously explained by DN. When implementing SPC, we incorporate the nonnegativity constraints for the response (i.e., **s** *≥* 0) so that the model response represents the firing rate, given that nonnegative sparse coding keeps the same expressiveness as standard sparse coding (Papyan et al., 2017).

#### 3.4.1 A homeostasis function for SPC

When using sparse coding to learn Gabor-like features that are similar to simple cells in V1 (Olshausen and Field, 1996, 1997), homeostasis functions were chosen to be of the particular forms given by

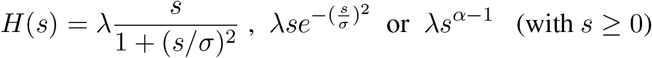

where *α* = 1 (Olshausen and Field, 1996, 1997; Zhu and Rozell, 2013; Lian et al., 2019). *H*(*s*) = *λs*^*α*−1^ with *α* = 2 is used by Rao and Ballard (1999), but the model does not generate Gabor-like receptive fields. When *H*(*s*) takes the form of *λs*^*α*−1^ (with *s ≥* 0), from our simulation, we find that it is necessary to choose *α* in the range 1 *≤ α <* 2 to enable the model to learn Gabor-like receptive fields, as will be shown in Section 3.4.2 and illustrated in Fig. 5.

**Figure 5:**
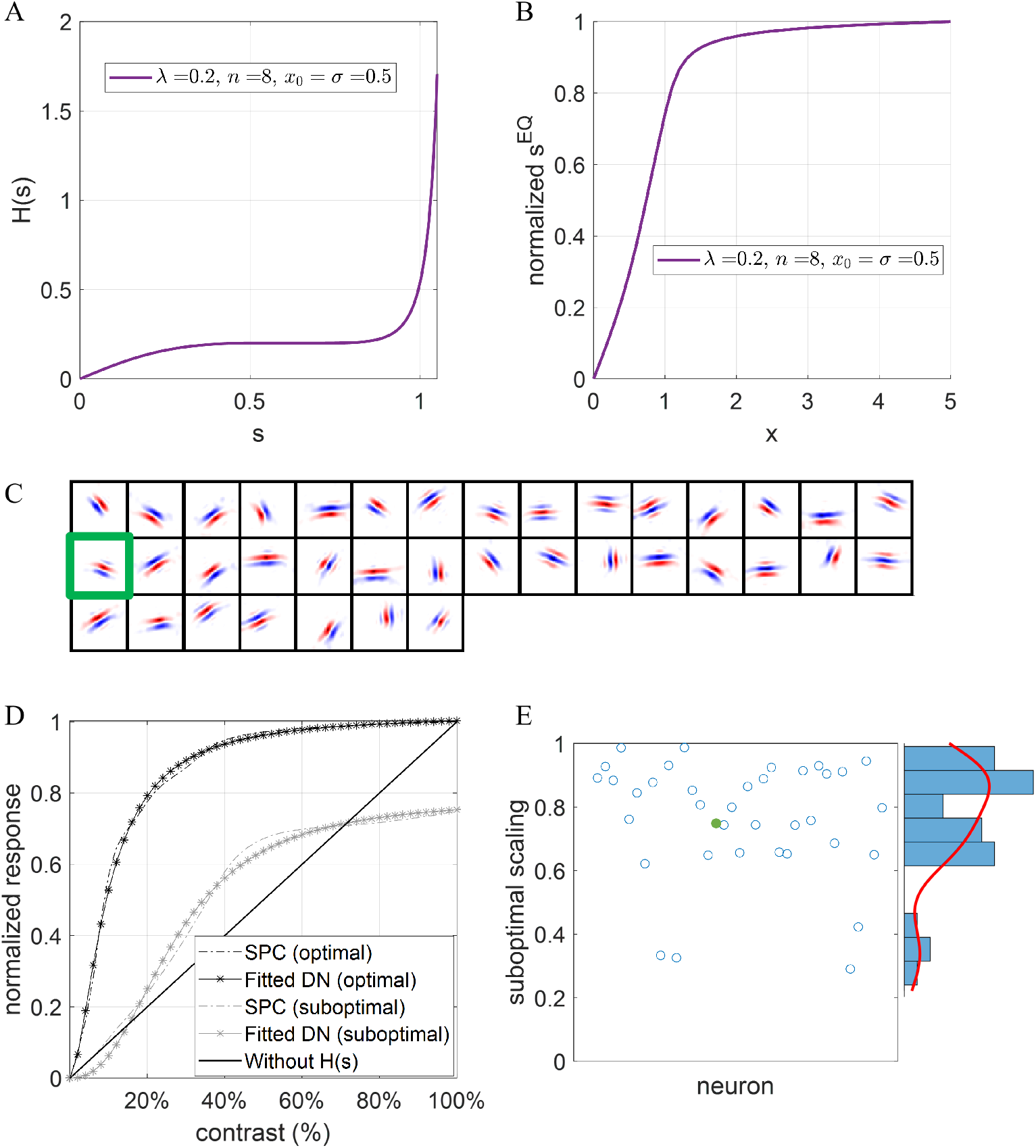
SPC with a proposed homeostasis function can learn simple cells as well as account for response saturation. (A) The proposed homeostasis function described by Eq. 22. In this example, parameters are chosen as follow: *λ* = 0.5, *x*_0_ = 0.5, *σ* = 0.5 and *n* = 4. (B) The corresponding analytical response vs. input curve by computing the intersect (*s*^EQ^) between *Y* (*s*) and *H*(*s*) in Fig. 3. (C) Sparse/Predictive coding (SPC) model with this homeostasis function can learn Gabor-like features. Receptive fields of 37 model neurons whose receptive fields are located in the central regions and well within the visual visual field are shown and each square is a 16 *×* 16 image that shows the corresponding learnt feature. The green box is the receptive field of the learnt neuron whose contrast-response curve is displayed in D. (D) ‘SPC (optimal/suboptimal)’ represents normalized response of the model neuron (green box in C) of the SPC model when sinusoidal gratings with the optimal/suboptimal orientation are presented. ‘Fitted DN’ shows the closest DN fit by Eq. 16. ‘Without *H*(*s*)’ represents the response vs. contrast curve when the homeostasis function is removed after learning. (E) The scatter plot and histogram of the ratio between optimal and suboptimal responses (suboptimal scaling) for these 37 model neurons shown in C. Green dot represents the neuron in D whose suboptimal scaling value is close to the mean of the distribution in E.

After examining the homeostasis functions that can learn Gabor-like receptive fields for SPC models, one thing in common is that the homeostasis function is concave starting from 0. In order to introduce response saturation into the homeostasis function, the homeostasis function should approach infinity as the input increases, and a convex curvature of the function is necessary. Therefore, we propose a homeostasis function for the SPC model described as follows:

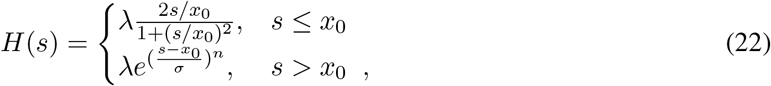

where *x*_0_ *>* 0 is the turning point at which the homeostatic function, *H*(*s*), becomes convex instead of concave, *σ* determines the steepness of *H*(*s*) when *s* is larger than *x*_0_, and *n* is the index that determines the steepness of the curve when *s* is large. An example of *H*(*s*) given by Eq. 22 is shown in Fig. 5A. Note that this homeostasis function has a similar shape to the corresponding homeostasis function of DN in the single-neuron scenario (Eq. 19 and Fig. 4B).

The proposed homeostasis function, *H*(*s*), in Eq. 22 has the following properties: (1) it is concave when *s < x*_0_, which facilitates the emergence of features after learning; (2) it is convex when *s > x*_0_; (3) the derivative at *s* = *x*_0_ is 0 such that the transition between concave and convex is smooth and continuous; (4) it approaches large values fairly quickly when *s* is large, which leads to response saturation. Note that the homeostasis function proposed here is designed to precisely specify the function shape in different regions of the neural activity to further investigate how different curvatures of the function would affect response nonlinearities.

Assuming **A**^*T*^ **A** = 1 and **A**^*T*^ **x** = *x* (where *x* represents the contrast or input amplitude), we can compute the analytical response vs. contrast curve by continuously changing *x* while numerically computing the intersect between *Y* (**s**) and *H*(**s**) in Fig. 3. The corresponding analytical response vs. contrast curve of *H*(*s*) in Fig. 5A is shown in Fig. 5B. Though **A**^*T*^ **A** = 1 does not always hold, this analytical response curve can still provide insights into the response linearity of the model because **A**^*T*^ **A** are normally very close to identity matrix after learning in practice.

#### 3.4.2 The model can learn Gabor-like simple cells while displaying contrast saturation

The SPC model can learn Gabor-like simple cells when a homeostasis function, *H*(*s*), as described in Fig. 5A. Fig. 5C shows receptive fields of 37 model neurons learnt by this SPC model with the above homeostasis that meet the requirements specified in Methods 2.4.2. The pattern of responses illustrates that the model learns Gabor-like features similar to those of simple cells in V1 and similar to previous modelling studies based on sparse coding (Olshausen and Field, 1996, 1997; Hoyer, 2003; Zylberberg et al., 2011; Zhu and Rozell, 2013; Lian et al., 2019). This shows that SPC with a suitable homeostasis function can learn features and it is not confined to a specific function as used in previous models, which is also validated by a recent study (Calatroni et al., 2023).

After learning, the response vs. contrast curve was examined to investigate the response nonlinearity of the model using. The response vs. contrast curve is then fitted by the equation of single-neuron DN (Eq. 16), similar to previous studies (Heeger, 1992; Busse et al., 2009). Fig. 5D shows that one example of learnt model cells displays contrast saturation for both optimal and suboptimal sinusoidal gratings and the response nonlinearity can be well fitted by DN. The ratio between optimal and suboptimal responses (defined as suboptimal scaling) is 0.75, indicating a lower response when sinusoidal gratings with non-preferred orientation are presented. Furthermore, the response curve is qualitatively similar to the analytical response curve as shown in Fig. 5B. If homeostasis function *H*(*s*) is removed after learning, the response vs. contrast curve becomes linear that shows no contrast saturation. Fig. 5E shows the scatter plot and histogram of suboptimal scaling values for all these 37 model neurons, which demonstrates a diversity of the extent to which model neuron response is reduced when suboptimal sinusoidal gratings are presented. Altogether, above results show that the proposed SPC model with *H*(*s*) in Fig. 5A can learn simple cell receptive fields from natural images while displaying response saturation at the same time.

We further train the proposed SPC model with another 3 different homeostasis functions as shown in Fig. 6A with their corresponding analytical response curve in Fig. 6B. For all these four model datasets, we plot the response vs. contrast curves for each model cell whose receptive field is well within the visual field (Methods 2.4.2). Fig. 6C shows the response vs. contrast curve for all four model datasets with different choices of homeostasis function *H*(*s*), similar to the distribution of values for experimentally measured cells shown in Fig. 6D. Moreover, different homeostasis functions *H*(*s*) can lead to diverse response nonlinearities and the analytical response curves (shown in Fig. 6B) give a decent approximation of response nonlinearities of model cells. Experimental data (Fig. 6D) has included cells that show no response saturation as well as those cells that saturate very quickly. More choices of the parameters in the proposed homeostasis function (Eq. 22) and wider family of homeostasis functions may account for more diversity of response nonlinearity.

**Figure 6:**
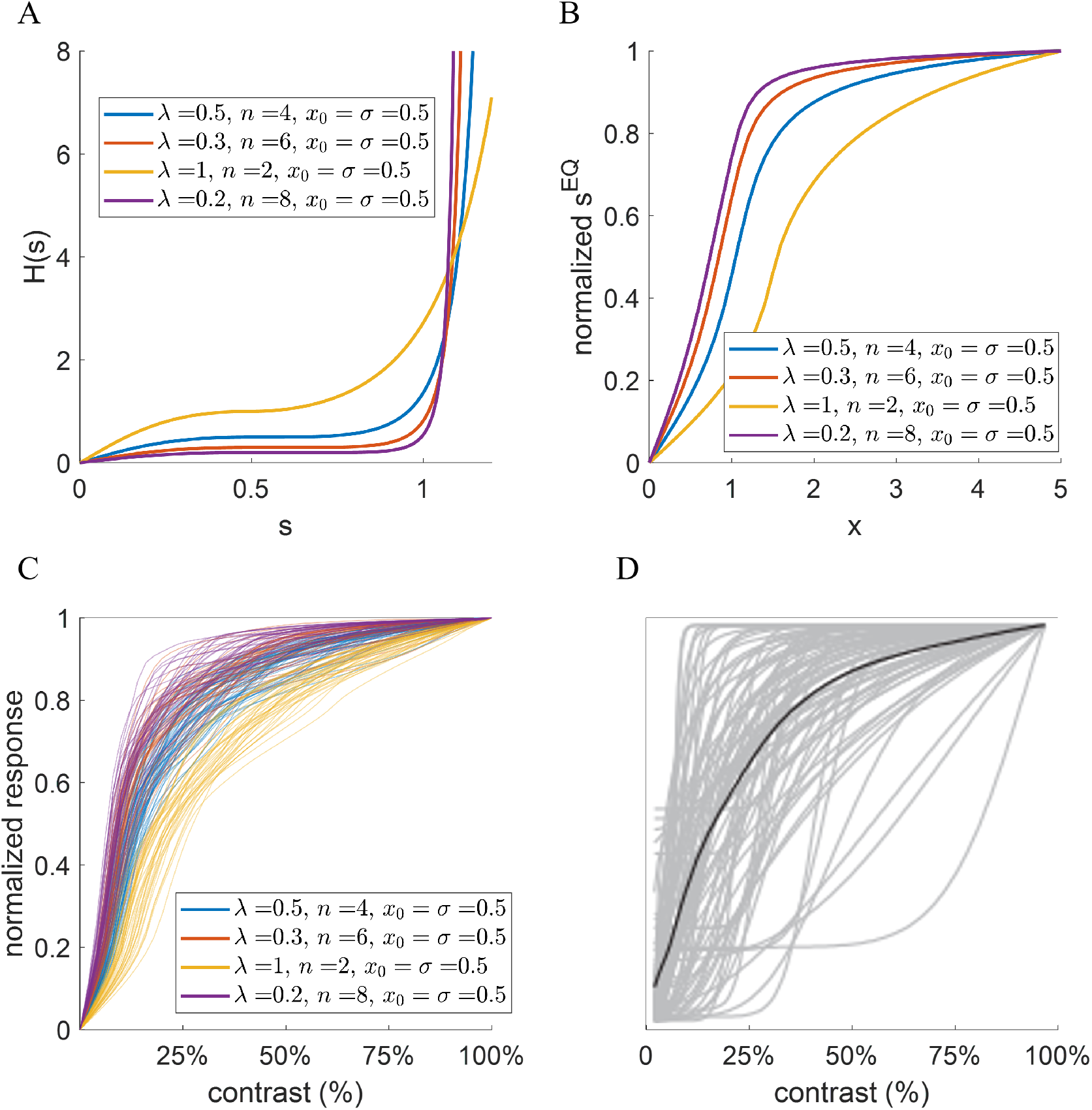
Sparse/Predictive coding (SPC) model with different homeostasis functions can generate diverse nonlinearities of response vs. contrast curve that are observed experimentally. (A) The proposed homeostasis function described by Eq. 22 with four different choices of parameter values. (B) The corresponding analytical response vs. input curve by computing the intersect (*s*^EQ^) between *Y* (*s*) and *H*(*s*) in Fig. 3. (C) Response vs. contrast curves for learnt simple cells whose receptive fields are centered well within the 16 *×* 16 visual field. The response is scaled to a maximum of 1 for each learnt cell. (D) Response vs. contrast curves for cells in cat V1 from an experimental study (adapted from (Busse et al., 2009)). The response is also scaled to a maximum of 1 for each cell. The black line represents the population average.

Overall, using the homeostasis function defined in Eq. 22, the SPC model possesses two important features: learning receptive field-like features and displaying nonlinearities similar to those observed experimentally. The simulations shown here, trained on natural images, demonstrate that the SPC model can learn Gabor-like simple cells as well as display contrast-saturation. Consequently, this SPC framework has great potential of describing and explaining how the neural circuits in the visual pathway possess nonlinear properties of the visual input that are observed experimentally.

## 4 Discussion

In this study, we aim to provide a framework for understanding sparse/predictive coding and divisive normalization (DN). Sparse coding and predictive coding can be identical in a two-layered neural network and unified as a sparse/predictive coding (SPC) framework whose model dynamics is determined by the predictive structure and a homeostasis function. We analytically showed that the SPC model can be identical to divisive normalization for a single-neuron scenario, namely that SPC with a homeostasis function can be derived from any divisive normalization function. The relationship between SPC and divisive normalization within a network with multiple neurons was also given. In a neural network model of V1, simulation results here show that SPC with a specific homeostasis function can learn Gabor-like simple cells, which gives similar receptive field properties to previous sparse and predictive coding model results (Olshausen and Field, 1996, 1997; Rao and Ballard, 1999), as well as displaying contrast saturation, which has previously been explained by DN (Heeger, 1992). In this framework, the homeostasis function, *H*(*·*), regulates the response of model units based on the current response level, its shape also leads to the emergence of sparse features.

The work presented here is also related with other previous studies. Rubin et al. (2015) demonstrate that a neural network with excitatory and inhibitory neurons, input-output power function, and fine-tuned recurrent connection weights can account for nonlinearities previously explained by DN. However, their work does not incorporate any learning ability of the network, nor the connection with sparse/predictive coding. Chalk et al. (2018) provides a framework of sparse and predictive coding from the perspective of information theory. However, their work does not consider divisive normalization and, unlike classical hierarchical predictive coding (Rao and Ballard, 1999), it addresses predictive coding in the temporal domain. The sparse/predictive coding model implemented here is similar to the study by Boutin et al. (2020), but their work does not consider the relationship with divisive normalization. Calatroni et al. (2023) has recently investigated the effects of different choices of sparsity function in the cost function of sparse coding that in essence is similar to changing the homeostasis function *H*(*s*) in this work, but their focus is not about how sparse coding can be connected with divisive normalization via the choice of homeostasis function. Our work unifies sparse coding, predictive coding and divisive normalization from first principles, showing how the non-linear response properties arise through learning (i.e., synaptic plasticity).

Winner-take-all, the mechanism in which only the neuron with the highest firing rate of the neural network is allowed to fire, has been used in many studies of computational neuroscience. The beneficial property of winner-take-all is that it provides strong competition to the neural network, which assists with learning useful features from the input (Einhäuser et al., 2002; Rennó-Costa and Tort, 2017), and can be accomplished by lateral inhibition within the network (Coultrip et al., 1992). The SPC framework proposed here is closely related to the winner-take-all mechanism; indeed winner-take-all can be viewed as an extreme sparse coding model where only a single neuron or small number of neurons fire, which can be implemented in the SPC model with a strong sparsity constraint. For example, when the homeostasis function, *H*(*·*), is a constant function, and the value of the constant determines the fraction of active neurons in the network.

Homeostasis is a well-established and prominent feature of the brain’s neural circuits, in which the brain employs a balancing mechanism to regulate neuronal excitatory and inhibitory activity (Turrigiano, 2011). Previously, Perrinet (2010, 2019) has incorporated synaptic homeostasis (weights scaling) into sparse coding and showed that V1 simple cells can emerge via an unsupervised learning process, similar to this work. However, in the SPC framework proposed in this study, the homeostasis function in the model dynamics, Eq. 10, regulates the response of each single neuron in a dynamic fashion, analogous to the homeostatic regulation of neuronal firing of biological neural circuits (Turrigiano, 2011). In the learning step of the SPC framework, connection weights are normalized after each update, which is equivalent to the effect of homeostatic synaptic rescaling (Turrigiano, 2011). The homeostasis function in the SPC model, however, has its origin in the mathematical derivation of the original sparse and predictive coding model, and its homeostatic role in regulating the neural circuits is somewhat fortuitous. The combined SPC-DN framework proposed in this study combines Hebbian and homeostatic learning and may serve as a more general framework for the understanding of nonlinear neural response properties and neural learning in brain circuits (Turrigiano, 2017).

The original predictive coding formulation (Rao and Ballard, 1999) is predictive in a hierarchical fashion, i.e., the model predicts the input in lower levels of the hierarchy from the activity in the higher levels, rather than being predicting in a temporal fashion, i.e., predicting future input. However, the input to the brain typically changes over time and the brain is constantly making predictions of the future on the basis of previous sensory stimuli and neural activity. In this paper, the visual input to the model is considered as static, consisting of a sequence of stationary images. However, this framework can also be used to investigate the problem of temporal prediction by incorporating synaptic delay between adjacent layers of the network hierarchy, as done in recent studies investigating predictive coding with temporal alignment (Hogendoorn and Burkitt, 2019; Burkitt and Hogendoorn, 2021).

The SPC framework proposed in this paper unifies sparse coding and predictive coding, and bridges its connection with divisive normalization. A predictive structure with feedforward input and feedback from higher layers preserves the essence of predictive coding and the model’s dynamics, given in Eq. 10, implements the dynamics of the neural activity in the network. However, a more fully biological implementation of this model would still require numerous additional features. First, the network would need to obey Dale’s Law by having separate populations of excitatory and inhibitory neurons. Second, spiking neurons that learn using spike-based synaptic plasticity, such as spike-timing-dependant-plasticity, would need to be used rather than the rate-based neurons typically used. Third, a more biological realization of the homeostasis function, which could differ for different neural populations, would be required. In addition, the choice of time constants of the dynamic system, synaptic delay between subsequent layers of the hierarchy, and many other parameters would need to be matched with experimental data for the particular neural circuit being modelled. Though some of these aspects have been considered in previous studies (Denève and Machens, 2016; Lian et al., 2019), much research still remains to be undertaken.

Given the success of divisive normalization in modelling data of multisensory integration (Ohshiro et al., 2011, 2017), this SPC framework might be able to uncover how the brain learns to integrate input from multiple sensory systems.

## Supporting Information

**Appendix**. This document has some further analysis of the work.

